# Toxicity of MAPT 4R RNA Contributes to Motor Neuron Degeneration in ALS

**DOI:** 10.64898/2026.09.22.753400

**Authors:** Hamish Crerar, Michele Stasi, Koustav Pal, Sabrina Pia Nuccio, Marija Petrić Howe, Giulia Manferrari, Yiran Wang, Benjamin Clarke, Marc-David Ruepp, Marco di Antonio, Rickie Patani

## Abstract

MAPT (Tau) dysregulation is implicated in several neurodegenerative diseases, but its contribution to amyotrophic lateral sclerosis (ALS) is poorly understood. Here we show that mRNA isoforms encoding 4-repeat (4R) Tau are upregulated and cytoplasmically enriched in iPSC-derived motor neurons (MNs) from VCP-mutant and sporadic ALS, without a corresponding change in Tau protein. Using splice-switching antisense oligonucleotides and isoform-specific siRNAs, we find that enhanced 4R expression reduces MN viability, whereas its selective knockdown improves survival, with kinetics more consistent with an RNA-intrinsic effect than altered protein synthesis. Exon 10-containing MAPT RNA shows increased predicted secondary structure, self-association and altered Tau biocondensation in vitro. In post-mortem ALS cervical spinal cord, increased relative exon 10 usage is associated with a higher-risk clinical phenotype and shorter disease duration These findings identify an isoform-specific contribution of MAPT to MN vulnerability in ALS and nominate 4R MAPT RNA as a therapeutic target.

## Introduction

Amyotrophic lateral sclerosis (ALS) is a fatal neurodegenerative disease characterised by progressive loss of upper and lower motor neurons. Despite significant advances in the genetic and cellular underpinnings of ALS, the mechanisms by which disruption of nuclear RNA metabolism is translated into selective motor neuron degeneration remain incompletely understood. A hallmark of ALS pathology across both familial and sporadic cases is the mislocalisation and eventual nuclear depletion of RNA-binding proteins (RBPs), such as TDP-43 ^1^, FUS ^2^, SFPQ^3^, ELAVL3 ^4^ and others ^5^. These RBPs regulate diverse aspects of RNA metabolism, including splicing, polyadenylation and transcript stability^6–9^. Disruption of their nuclear functions has been shown to alter the expression and processing of numerous neuronal RNAs ^10,11,8,12^, although the identity and functional impact of critical downstream targets remain to be fully elucidated.

Accumulating evidence implicates the dysregulation of microtubule-associated protein tau (MAPT) as a potentially important output of RBP dysfunction ^13,14^. *MAPT* is a highly regulated, alternatively spliced gene which produces isoforms containing either three or four microtubule-binding repeats (3R or 4R), with the balance of these isoforms tightly regulated during development and in the adult nervous system ^15^. Dysregulation of *MAPT* splicing is a defining feature of several primary tauopathies, including frontotemporal dementia (FTD), progressive supranuclear palsy, and corticobasal degeneration. While ALS is not traditionally classified as a tauopathy, emerging evidence suggests that tau dysfunction may also contribute to pathology within the ALS–FTD spectrum ^16–19^. Clinical cases can include both pure ALS, pure FTD and ALS-FTD. Recent studies have revealed that altered levels of Tau in plasma extracellular vesicles are diagnostic of ALS-FTD spectrum disorders ^20^ Elevated levels of phosphorylated tau (pTau) have been detected in post-mortem brain and spinal cord tissue, relative to the cerebellum, from patients with ALS ^21^. Functionally, studies in *VCP* mutant models of ALS–FTD have shown altered tau splicing, aggregation, and increased tau phosphorylation ^22^, supporting a role for tau as a modulator of neuronal health in this context.

Mechanistically, regulation of MAPT exon 10 has been directly linked to ALS-associated RBPs. Both FUS and SFPQ have been demonstrated to directly regulate splicing of *MAPT* pre-mRNA at exon 10, the site that determines the production of 4R versus 3R isoforms ^13^. Spatial dislocation of FUS and SFPQ within the nucleus leads to increased inclusion of exon 10 and a shift toward 4R MAPT expression. Indeed, the frontal cortex of ALS-FTD patients display increased levels of exon 10 relative to control ^23^. This raises the possibility that mislocalisation and dysregulation of FUS and SFPQ in ALS may drive isoform-specific *MAPT* dysregulation, with potential consequences for neuronal function and viability. Moreover, TDP-43 has been shown to directly regulate Tau at the level of mRNA stability, binding to UG-rich elements within the 3’UTR of *MAPT* transcripts and promoting their degradation. Notably, cytoplasmic mislocalisation of TDP-43 is associated with increased tau expression in mouse brains^14^.

Despite growing evidence linking RBP dysfunction to altered *MAPT* splicing, the downstream consequences of aberrant exon 10 inclusion remain poorly understood. The majority of existing models have assumed that pathological effects are induced through increased production of 4R protein. However, whether exon 10-containing MAPT transcripts themselves possess biological properties that contribute directly to neuronal dysfunction has not been investigated. Given the emerging acceptance that RNA molecules contribute to cellular function beyond their roles as templates for protein synthesis ^24–26^, we hypothesised that exon 10 inclusion may generate a *MAPT* transcript with distinct properties that directly contribute towards motor neuron degeneration.

Here, we directly address a critical knowledge void in ALS pathogenesis by revealing an increase in *MAPT* transcript isoforms encoding 4R Tau in human sporadic ALS and *VCP* mutant iPSC-derived motor neurons, with a cytoplasmic enrichment. Although total Tau protein abundance remains unchanged, exon 10-containing transcripts promote motor neuron degeneration and exhibit distinct structural properties and protein interaction dynamics. Together our findings suggest that aberrant *MAPT* splicing generates an RNA isoform with pathogenic properties beyond its canonical role in protein production, identifying RNA-mediated dysfunction as a previously unrecognised consequence of *MAPT* dysregulation in ALS.

## Results

### ALS Human Motor Neurons Exhibit Dysregulation of MAPT expression

We first analysed *MAPT* transcript levels across four developmentally defined time points across motor neuron (MN) lineage restriction, including induced pluripotent stem cells (iPSCs; DIV=0), motor neural (ventral spinal) precursors (mNPCs; DIV=14) and terminally differentiated MNs (iPSC-MNs) at two time points (DIV=24 and 35). We observed a progressive and significant upregulation of *MAPT* expression during differentiation, however this was not linear. *MAPT* levels rose modestly between iPSCs to neural precursors, followed by marked and significant increase occurring upon terminal differentiation into iPSC-MNs, with a more modest increase following further maturation (Extended Data Figure 1A).

When comparing terminally differentiated iPSC-MNs derived from healthy controls (CTRL), sporadic ALS (sALS), and VCP-mutant (VCP^mu^) iPSCs, distinct patterns emerged. In sALS MNs, 4-repeat (4R) *MAPT* isoform expression was significantly increased, while total MAPT transcript levels remained unchanged (Figure 1A), suggesting a direct shift in isoform expression. In contrast, *VCP^mu^* MNs showed a significant increase in both total *MAPT* and *4R MAPT* isoform expression relative to controls (Figure 1B), indicating that *MAPT* dysregulation in ALS may arise through multiple mechanisms but importantly, it converges on elevated exon-10 inclusion.

**Figure 1:**
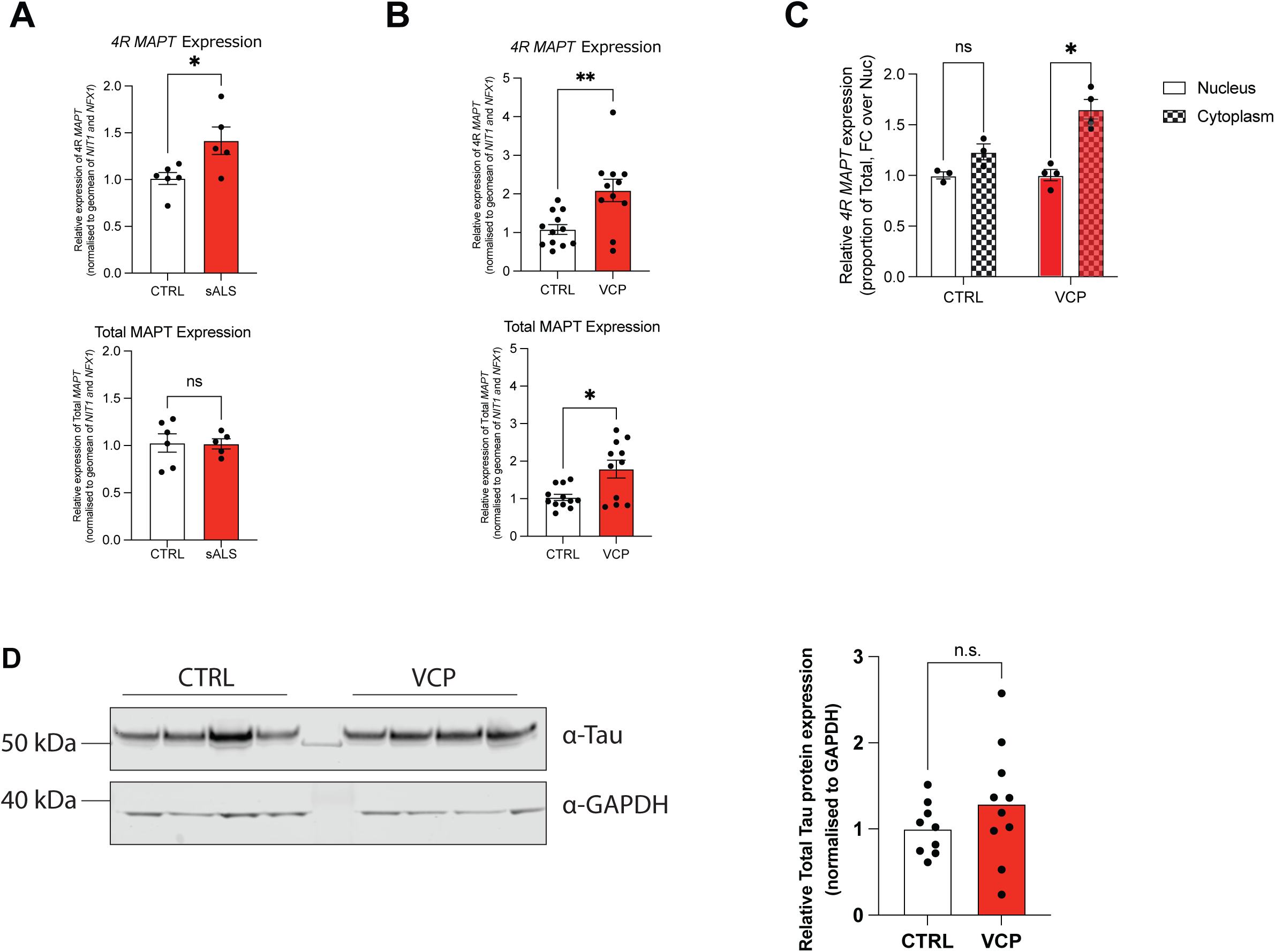
**A** qRT-PCR analysis of expression of exon 10 containing *4R MAPT* (*Top*) or Total *MAPT* (Bottom) in CTRL or sALS iPSC-MNs. Values are normalised over the geometric mean of housekeeper genes *NIT1* and *NFX1* and expressed as fold change over CTRL. These were chosen as we have previously demonstrated their levels to be highly stable across patient and control conditions^35^. Data points indicate individual patient lines. Unpaired t-Test with Welch’s correction with significance placed at P<0.05. **B** as for (A) with CTRL and *VCP^mu^* iPSC-MNs, data points indicate individual patient lines from a minimum of three inductions. **C** MAPT exon 10 expression levels in *VCP^mu^* ALS iPSC-MN nuclear and cytosolic fractions. Data normalized by expression of *MAPT* exon 10 over total levels of *MAPT* expression. Multiple unpaired t-Test with Holm-Sidak multiple comparison with significance placed at P<0.05. **D (**Left) Representative western blots showing total Tau, and GAPDH loading control in CTRL and *VCP^mu^* iPSC-MNs. (Right). Densitometry of blots. Values are normalised over GAPDH levels and expressed as fold change over CTRL. Data points indicate individual iPSC lines from different patient donors, from three separate inductions. Unpaired t-Test with Welch’s correction with significance placed at P<0.05. ns = not significant, * =P<0.05 **=P<0.001

We have previously demonstrated that a defining hallmark of ALS is the aberrant localisation of transcripts between the nuclear and cytoplasmic subcellular compartments^2,3^, which may escape detection in whole-cell analyses. To address the hypothesis that the *4R MAPT* isoform exhibits subcellular mislocalisation, we performed nuclear–cytoplasmic fractionation followed by isoform-specific quantification by qPCR. This analysis revealed a significant enrichment of *4R MAPT* RNA in the cytoplasm of *VCP^mu^* MNs, an effect not observed in control neurons (Figure 1C).

Interestingly, despite these transcriptional changes, western blotting revealed no significant difference in total Tau protein abundance between VCP mutant and control iPSC-MNs (Figure 1D). Furthermore, 4R Tau protein could not be reliably detected by western blotting under these differentiation conditions, consistent with previous studies reporting that iPSC-derived motor neurons predominantly retain a developmentally immature Tau isoform profile following relatively short differentiation paradigms^27^. Together, these data suggest that increased expression of exon 10 containing *MAPT* transcripts does not result in detectable changes in Tau protein abundance in iPSC-MNs.

### Modulating MAPT Isoform Usage Alters Motor Neuron Survival

We next sought to investigate if the alterations we observe in *MAPT* isoform expression play a causative role in previously demonstrated *VCP^mu^* MN survival phenotypes ^28^. We employed a longitudinal live-cell imaging approach, which has been shown to have higher sensitivity than cross sectional approaches ^28,29^, to assess levels of iPSC-MN death. In the untreated condition, we observed significantly increased death in *VCP^mu^*iPSC-MNs, as early as 24h into our differentiation paradigm (Figure 2A). The increased iPSC-MN death remains significant at each timepoint assayed (24 hour intervals for 5 days post differentiation). We next selectively manipulated *MAPT* isoform expression utilising transfection of siRNA and antisense oligonucleotide (ASO) approaches to evaluate necessity and sufficiency. A splice-switching ASO designed to promote exon 10 inclusion ^30^ increased *4R MAPT* levels without altering total transcript levels (Extended Data Figure 2A). Conversely, an siRNA targeting exon 10 of *MAPT* selectively reduced 4R isoform expression without affecting total *MAPT* transcript levels, while a pan-MAPT siRNA reduced both (Extended Data Figure 2B). At the protein level, pan-MAPT siRNA led to a clear reduction in total Tau, as assessed by western blot using a pan-Tau antibody, whereas 4R-specific siRNA had no detectable effect on total Tau levels (Extended Data Figure 2C), consistent with its isoform-specific targeting and low levels of 4R Tau protein. Treatment with the ASO promoting exon 10 inclusion increased cell death in CTRL iPSC-MN (Figure 2B) and exacerbated the pre-existing survival phenotype in *VCP^mu^* iPSC-MN (Figure 2C). This suggests that increased expression of exon 10 containing *MAPT* transcripts is sufficient to cause iPSC-MN death. Conversely, knockdown of *4R MAPT* using isoform-specific siRNA significantly rescued the survival phenotype in *VCP^mu^* MNs (Figure 2D). Notably, total *MAPT* knockdown failed to confer a similar protective effect, despite effectively reducing total Tau protein abundance (Extended Data Figure 2C). These findings indicate that motor neuron survival is influenced by the selective abundance of exon 10-containing MAPT transcripts rather than overall MAPT expression.

**Figure 2:**
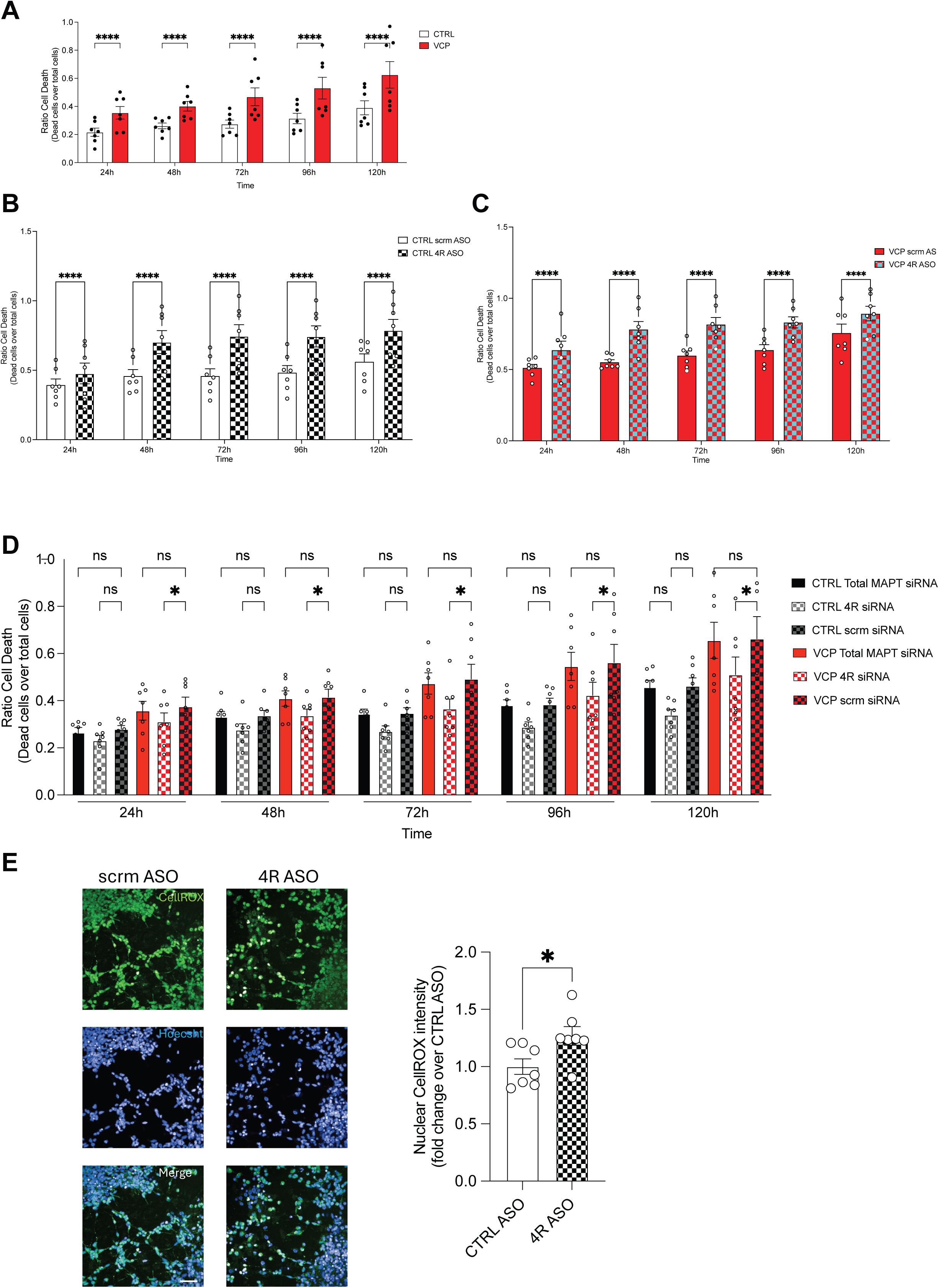
**A** Longitudinal live-cell imaging of CTRL and *VCP^mu^* iPSC-MNs over 5 days post differentiation was performed using the Incucyte live cell imaging and analysis system. Data points indicate individual lines from two inductions with two experimental repeats. Ordinary Two-Way ANOVA with Tukey’ multiple comparisons test and significance placed at P<0.05. **B.** Longitudinal live-cell imaging of CTRL iPSC-MNs transfected with ASO promoting exon 10 inclusion (4R ASO) ^30^ or scrambled control and imaged for 5 days post transfection. Data points indicate individual lines from two independent experimental repeats. Ordinary Two-Way ANOVA with Tukey’ multiple comparisons test and significance placed at P<0.05. **C** As (B) but with VCP MNs. **D** Longitudinal live-cell imaging of CTRL and *VCP^mu^* iPSC-MNs transfected with an siRNA targeting *MAPT* exon 10 selectively (*4R MAPT*), all *MAPT* isoforms (total *MAPT*) or a scrambled control and imaged for 5 days post transfection. Data points indicate individual lines from two independent experimental repeats. Ordinary Two-Way ANOVA with Tukey’ multiple comparisons test and significance placed at P<0.05. **E.** (Left) Representative fluorescence images of *VCP^mu^*iPSC-MNs transfected as indicated and stained with CellROX Green (green) to visualize overall ROS generation and Hoecht (blue). Scale bar, 50 μm. (Right) Quantification of nuclear CellROX Green intensity. Graphs plotted as fold change over average CTRL ASO transfected condition per technical replicate. Unpaired t-Test with Welch’s correction with significance placed at P<0.05. Data points indicate individual transfected lines from two independent experimental repeats. ns = not significant, * =P<0.05 ***=P<0.0001

Given the known role of SFPQ and FUS in regulating *MAPT*, and their previously demonstrated mislocalisation in ALS MNs ^2,3,5^, we next sought to determine whether changes in *MAPT* isoform use and subsequent phenotypes were upstream or downstream of RBP localisation changes. We assessed the localisation of FUS and SFPQ following siRNA and ASO treatment. Neither intervention affected the nuclear mislocalisation phenotype in iPSC-MNs (Extended Data Figure 2D), suggesting that altered *MAPT* isoform expression occurs downstream of or in parallel to RBP mislocalisation.

To further investigate the mechanisms underlying altered motor neuron survival following manipulation of MAPT isoform expression, we assessed oxidative stress using the CellROX assay. Previous work from our lab has shown that reactive oxygen species (ROS) are increased in VCPmu MNs ^28^ and alterations in Tau function have previously been implicated in oxidative stress pathways in neurons ^31^. Upon oxidation, CellROX undergoes oxidation-dependent nuclear translocation and binds DNA, providing a quantitative measure of intracellular oxidative stress. Consistent with the reduced survival observed following exon 10 inclusion, treatment with the 4R-promoting ASO resulted in a significant increase in nuclear CellROX intensity compared with scrambled ASO-treated controls (Figure 2E), indicating elevated oxidative stress following increased expression of exon 10-containing MAPT transcripts.

### Exon 10 modulates MAPT Tau condensation properties

To determine whether mislocalisation of SFPQ alters its association with *MAPT* RNA, we first analysed SFPQ iiCLIP data from (whole cell) iPSC-MNs, which revealed that SFPQ binding at MAPT exon 10, clearly detected in control neurons, was noticeably reduced in VCPmu lines (Figure 3A), consistent with expected loss in nuclear interaction. Because fractional iiCLIP is currently unfeasible technically from hiPSC-derived neurons given the amount of material required, we next performed cytoplasmic RNA immunoprecipitation (RIP) for SFPQ as a viable alternative to assess whether binding is redistributed rather than simply lost. This revealed an enrichment of total *MAPT* RNA in the cytoplasmic SFPQ pulldown for VCPmu MNs compared with CTRL iPSC-MNs, while pull-down of a validated SFPQ target transcripts *BCL2L2* and *LMNB2* ^32^, remained unchanged (Figure 3B). These results suggest that increased cytoplasmic association between SFPQ and *MAPT* RNA is specific and may reflect a pathological interaction arising from altered subcellular localisation of both the RBP and its RNA target.

**Figure 3.**
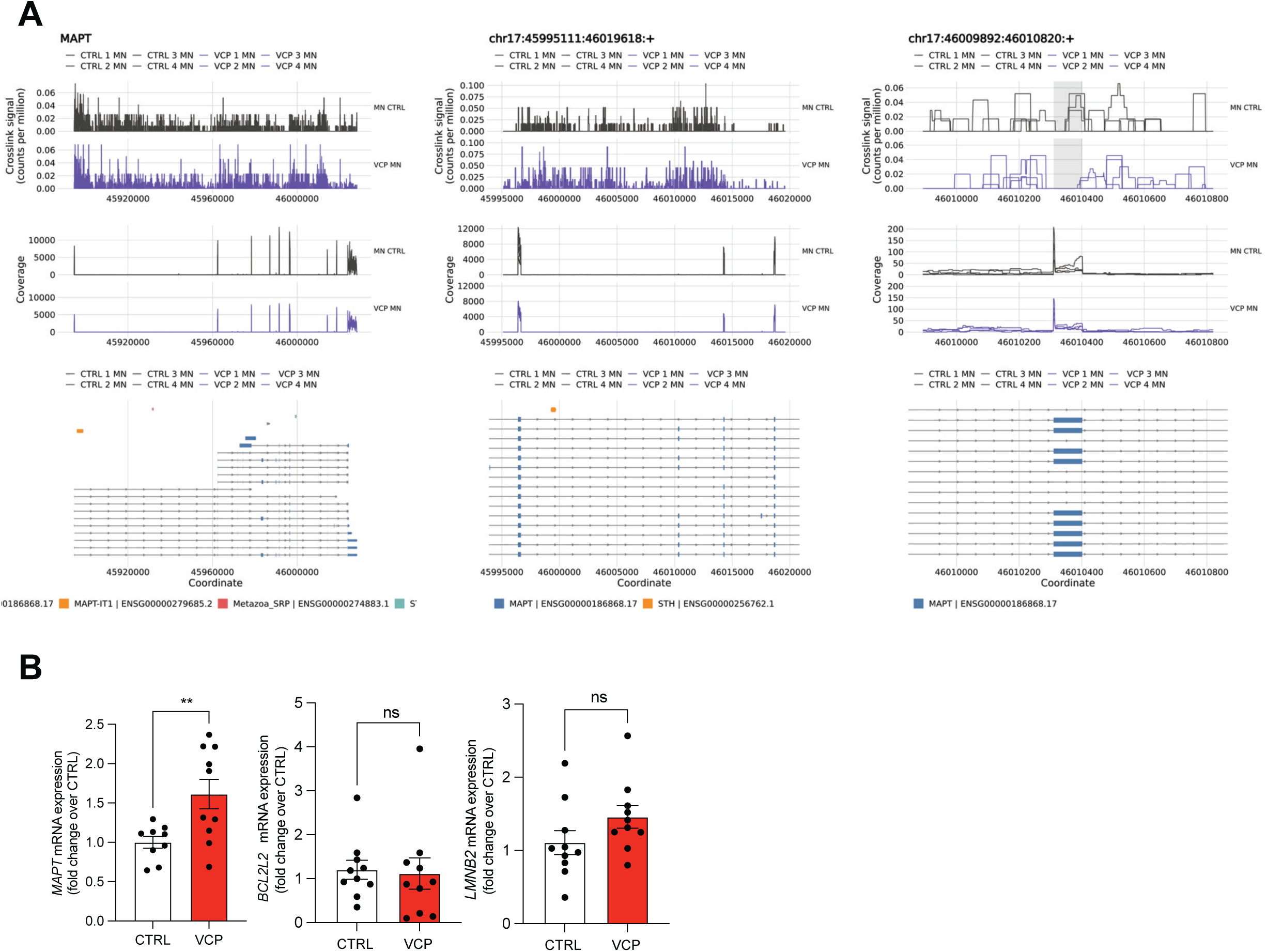
**A** Representative iiCLIP tracks generated with clipplotr showing SFPQ binding across the MAPT locus in control motor neurons (MN CTRL; black) and VCP mutant MN (VCP MN; purple). Top tracks show iiCLIP crosslinking signal normalised by library size; middle tracks show RNA-seq coverage. iiCLIP signal was smoothed using a 100-nt rolling mean. The bottom track shows MAPT transcript annotation. Left: overview of the MAPT gene. Centre: zoomed view of introns 9–10. Right: detailed view of SFPQ binding sites surrounding MAPT exon 10. **B** RNA immunoprecipitation (RIP) performed on the cytoplasmic lysates from control and *VCP^mu^* iPSC-MNs using antibodies for SFPQ or normal IgG as negative control. Levels of associated *MAPT*, *BCL2L2* or *LMNB2* expression were normalised by total input. Graphs plotted as fold change over average CTRL values. Data points indicate individual lines from three experimental repeats on three separate inductions. Unpaired T-Test with Welch’s Correction and significance placed at P<0.05. ns = not significant, **=P<0.001

We reasoned that this altered interaction may arise from intrinsic properties of the exon 10-containing transcript itself. ViennaRNA modelling predicted that inclusion of exon 10 substantially favours the formation of intramolecular *MAPT* RNA (Figure 4A). The minimal free energy (MFE) model and centroid secondary structure (CSS) models were markedly similar when exon 10 was present, consistent with a more compact and stable conformation, and higher likelihood of forming secondary structures. Consistent with this, RNA multimerisation modelling under near-physiological ionic conditions ([RNA] = 5 µM; [NaCl] = 150 mM; 25 °C) predicted that exon 10-containing species form intermolecular duplexes and higher multimers more readily than exons 11 and 12, indicating an increased propensity for *intermolecular* RNA–RNA base pairing and higher-order assembly (Figure 4B).

**Figure 4.**
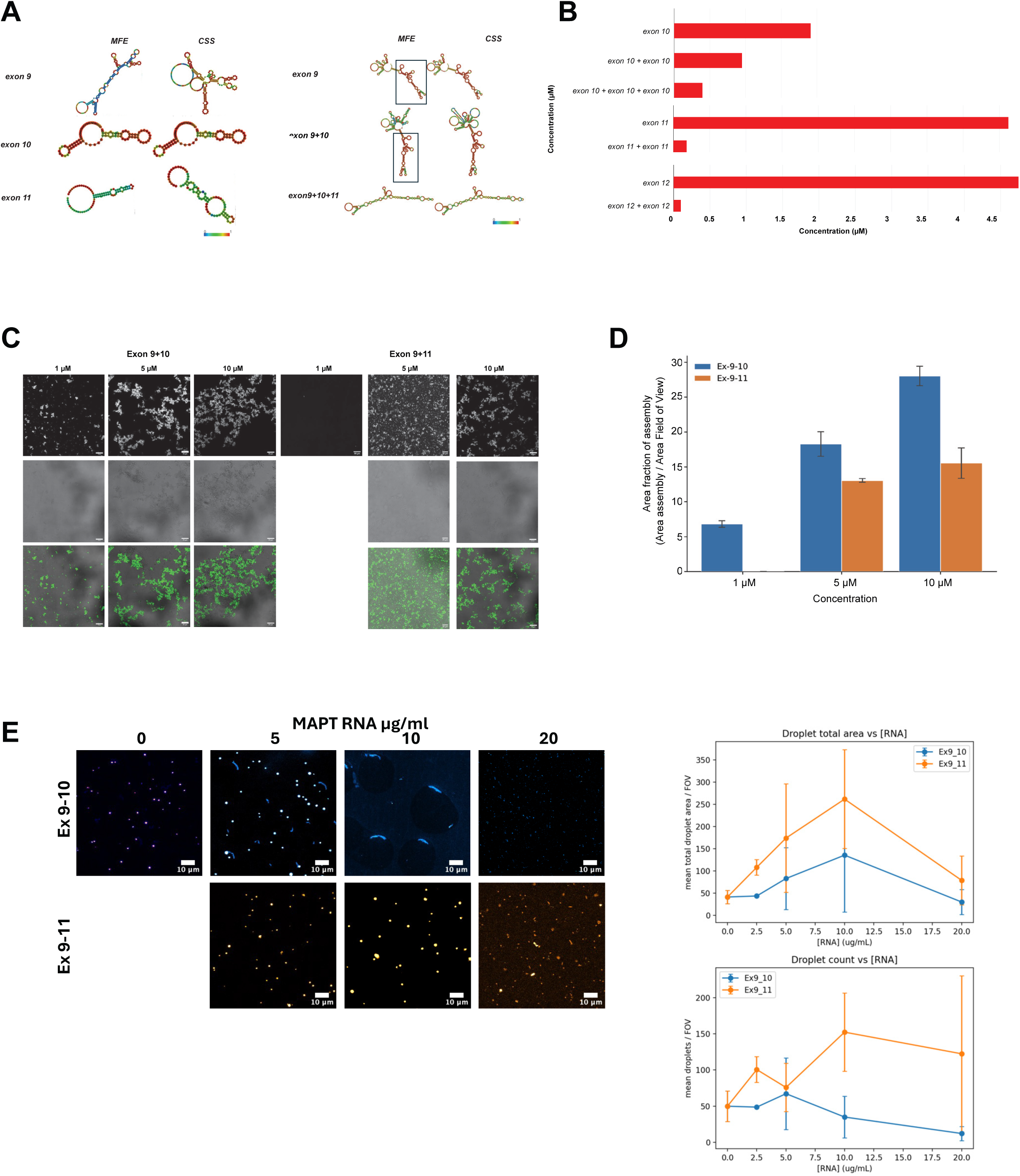
**A** ViennaRNA webserver simulation of RNA secondary structure of *MAPT* exons 9, 10 and 11, or exon 9+10, exon 9+10+11 and exon 9+11. No constraints were imposed on the simulation. Free energy of secondary structures accounting for transcript flexibility are calculated, outputting two likely possibilities; Minimum free energy (MFE), the most stable conformation, and Centroid Secondary Structure (CSS) **B** RNA multimerisation modelling under near-physiological conditions ([RNA] = 5 µM; [NaCl] = 150 mM; T = 25 °C) predicts that exon 10-containing species form intermolecular duplexes and triplexes more readily than exon 11 or 12. **C** Representative fluorescence micrographs of *in-vitro*-transcribed *MAPT* exon 9–10 and exon 9–11 RNA at indicated concentrations, incubated in 10 mM HEPES, 50 mM MgCl₂. **D** Semi-quantitative image analysis, measuring the area of assemblies after image segmentation over the area of the field of view, shows that exon 9-10 RNA forms larger clusters of aggregates at each concentration tested, supporting the self-association of RNA mediated by exon 10 (n=2, N=3). Error bar represents the standard deviation between replicates. **E** Representative fluorescence images of recombinant Tau incubated with in vitro-transcribed MAPT exon 9-10 or exon 9-11 RNA at increasing RNA concentrations (0–20 µg/mL) under macromolecular crowding conditions (n=2, N=4). Graphs show the number of Tau condensates (top) and total condensate area (bottom) as a function of RNA concentration. Scale bar represent 10 µm

Our experimental analysis supported these predictions: *in vitro*-transcribed exon 9-10 and exon 9-11 RNA fragments were subjected to RNase A digestion, which selectively cleaves single-stranded RNA. Therefore transcripts adopting more protected conformations are expected to exhibit increased resistance to digestion. Exon 9-10 RNA retained a larger proportion of full-length transcript relative to exon 9-11, indicating reduced susceptibility to single-stranded RNA cleavage (Extended Data Figure 3A). This observation is consistent with the increased structural stability and intermolecular interactions predicted by the ViennaRNA folding and multimerisation analyses. Similarly, exon 9-10 RNA underwent concentration-dependent self-association, with visible condensates forming at concentrations as low as 1 μM, whereas exon 9-11 RNA required at least 5 μM before comparable aggregation occurred (Figure 4C). Semi-quantitative image analysis, measuring the area occupied by segmented assemblies relative to the area of the field of view, showed greater assembly coverage for exon 9–10 RNA at each concentration tested (Figure 4D), supporting increased self-association mediated by exon 10.

Given the established role of RNA in regulating protein phase behaviour ^26,33^, we next investigated whether Tau directly interacts with MAPT RNA and whether exon 10-containing transcripts affect Tau condensation. Electrophoretic mobility shift assays (EMSA) confirmed direct binding between recombinant Tau protein and MAPT RNA *in vitro* (Extended Data Figure 3B). Under macromolecular crowding conditions, recombinant Tau readily formed condensates following incubation in the presence of PEG. Surprisingly, addition of exon 9-10 MAPT RNA resulted in a progressive loss of visible Tau condensates, whereas exon 9-11 RNA appeared to increase condensation and aggregation consistent with previous studies ^34^ (Figure 4E; Extended Data Figure 3C). Thus, although exon 10-containing MAPT RNA exhibits an increased propensity for self-association, it simultaneously reduces Tau condensation in vitro, demonstrating that exon 10 can differentially regulate both RNA-RNA and RNA-protein assembly.

Together, these findings strongly suggest that inclusion of exon 10 fundamentally alters the biophysical behaviour of MAPT RNA, increasing its structural stability and self-association while modifying Tau condensation dynamics. Together with the observed increase in cytoplasmic association with mislocalised SFPQ, these data suggest that exon 10 inclusion stabilises RNA structure and promotes RNA-protein interactions that may collectively contribute to RNA-driven toxicity in ALS motor neurons.

### MAPT relative Exon 10 usage is associated with clinical outcome in ALS

To explore whether aberrant *MAPT* exon 10 splicing extends beyond our iPSC experimental models and is associated with clinical disease, we analysed *MAPT* exon 10 expression levels in a cohort of ALS patients with matched clinical data from the New York Genome Center (NYGC) ALS Consortium. We investigated whether MAPT dysregulation was associated with clinical outcome. In cervical spinal cord, survival modelling incorporating MAPT expression alongside established clinical covariates, including age at symptom onset and genotypic sex, identified MAPT expression as a significant contributor to patient risk (Figure 5A). Stratification according to the model-derived risk score separated patients into groups with markedly different disease duration (HR = 2.255, log-rank P < 0.0001; Figure 5B). Patients within the high-risk group exhibited significantly lower total MAPT expression than those in the low-risk group (P = 0.009; Figure 5C).

**Figure 5.**
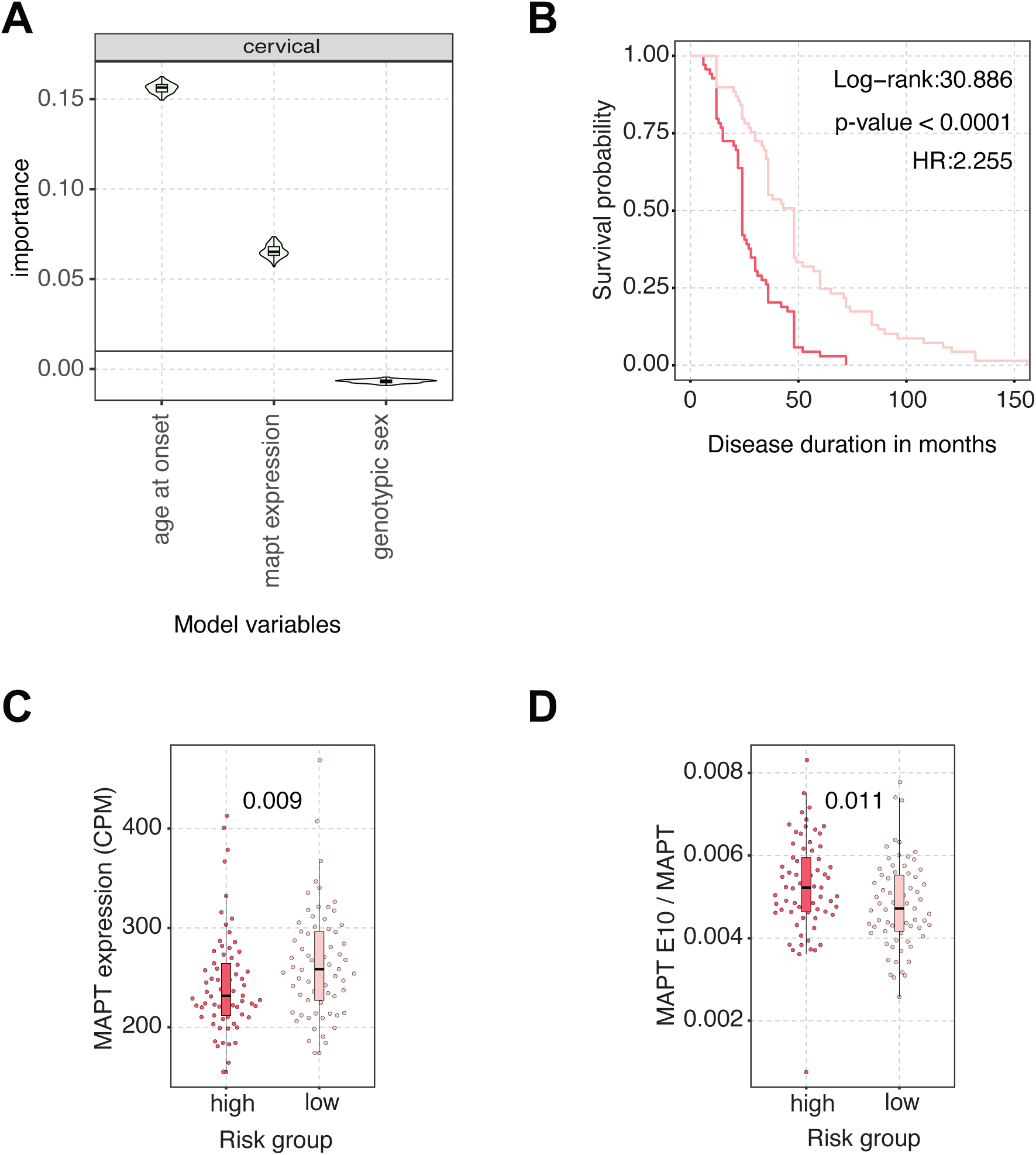
Investigating disease progression as a function of MAPT E10 usage in postmortem ALS cervical spinal cord samples. **A** Variable importance of covariates used for survival modelling. **B** Kaplan-Meier curves showing disease progression for ALS patients stratified into high and low risk groups. **C** MAPT expression in CPM units showcasing expression distribution in ALS patients stratified into high and low risk groups. **D** MAPT E10 usage in ALS patients stratified into high and low risk groups on the basis of MAPT expression.

Strikingly, despite this reduction in overall MAPT abundance, relative exon 10 usage was significantly increased in the high-risk group (P = 0.011; Figure 5D). This suggests that the clinical association is not simply reflective of decreased overall MAPT expression, but may instead involve altered relative usage of exon 10-containing MAPT transcripts.

Together with the increased exon 10-containing MAPT transcripts observed in ALS iPSC-derived motor neurons, these findings provide independent evidence that MAPT exon 10 dysregulation occurs in human ALS and that the enrichment of relative exon 10 usage in patients with poorer clinical outcome, despite reduced overall MAPT expression, supports selective dysregulation of exon 10-containing MAPT transcripts as a reproducible and clinically relevant molecular feature of ALS.

## Discussion

Our study reveals a novel and isoform-specific mechanism of *MAPT* dysregulation in ALS, thereby linking the previous discoveries of spatial dislocation^13,23^ and the mislocalisation of FUS and SFPQ^2,3,5^ to aberrant *MAPT* splicing and increased exon 10 inclusion. We identify that dysregulation of MAPT exon 10 is a convergent feature of ALS and that these transcripts contribute to motor neuron vulnerability in a manner likely beyond their canonical role as templates for Tau protein synthesis. Using patient-derived iPSC-MNs, we find that *MAPT* isoform regulation differs between ALS subtypes. In *VCP^mu^*lines, both *4R MAPT* and total *MAPT* transcript levels are increased, consistent with a broader dysregulation of *MAPT* expression. In contrast, sALS samples show a selective increase in *4R MAPT* without a change in total *MAPT* indicating isoform-specific regulation. These findings suggest distinct underlying mechanisms. A coordinated increase in total and 4R *MAPT* in *VCP^mu^* may reflect the global transcriptional changes consistent with this mutation ^3,5,35^, or an outcome of alterations in protein homeostasis due to altered function of VCP^36^. However, the selective increase in 4R *MAPT* in sALS points towards a dysfunction in splicing, shifting exon 10 inclusion without a change in overall transcript levels. These data suggest that although *MAPT* dysregulation is a shared phenomenon of ALS, the upstream drivers leading to this differ between sporadic and familial disease. Importantly, this pattern extends beyond iPSC models: analysis of large-scale NYGC post-mortem transcriptomic data Linked *MAPT* dysregulation to clinical outcome. Lower total *MAPT* expression was associated with a higher-risk clinical phenotype, yet these same patients exhibited significantly greater relative exon 10 usage. Together, these findings extend the relevance of MAPT biology beyond classical tauopathies and position exon 10-containing MAPT transcripts as clinically relevant molecular features of ALS.

A central finding of this study is that manipulation of *MAPT* exon 10 inclusion has clear effects on MN survival. Increasing exon 10 inclusion using splice-site switching ASOs reduced survival in control and exacerbated death in VCP^mu^ MNs. Conversely, selective depletion of exon 10 *MAPT* transcripts improves survival in VCP^mu^ , while total *MAPT* knockdown, despite reducing protein abundance, did not confer protection. These data together demonstrate that the survival phenotype is isoform-dependent rather than driven by total MAPT abundance, suggesting that reducing exon 10-containing transcripts may alleviate 4R MAPT-associated toxicity while preserving or favouring 3R isoforms, whereas pan-MAPT knockdown removes both and fails to restore the physiological 3R:4R balance. This is consistent with multiple studies ^37^ suggesting that the relative balance of 3R and 4R Tau isoforms, rather than their absolute abundance, is a critical determinant of neuronal homeostasis. This interpretation is further supported by the post-mortem analysis, in which patients with poorer clinical outcome exhibited increased relative exon 10 usage despite lower total *MAPT* expression, reinforcing the distinction between overall *MAPT* abundance and isoform-specific regulation. However, we did not directly demonstrate that 4R knockdown restores a protective 3R:4R stoichiometry, and this remains an important question for future work.

While tau protein pathology is well recognised in ALS–FTD spectrum disorders, particularly in cases with *MAPT* mutations or co-pathology, our data suggest that wild-type *MAPT* isoform dysregulation may also drive neurodegeneration through 4R RNA toxicity. The selective upregulation and cytoplasmic localisation of *4R MAPT* RNA in *VCP^mu^* neurons, coupled with the conspicuously rapid phenotypic rescue upon *4R MAPT* transcript knockdown, suggest a toxic gain of function at the RNA level. This is of particular note when the long half-life of Tau is considered, estimated to be over a week in iPSC MNs ^38^. Moreover, increased exon 10-containing *MAPT* RNA was not accompanied by a detectable increase in total Tau protein abundance in VCP^mu^ MNs, and 4R Tau protein remained below the detection threshold of our assays. Though these findings do not exclude a contribution from low-level or locally translated 4R Tau, they do raise the intriguing notion that the *MAPT* transcript itself may contribute directly to cellular toxicity, independent of its canonical role as a protein-coding gene.

Recent work provides mechanistic insight into how *MAPT* transcript and proteins might conspire to drive neuronal toxicity, with several studies highlighting a theme of RNA and RBP association with pathological Tau assemblies. For example, RNA-dependent 2N4R Tau fibrils contained associated 18S ribosomal RNA and were destabilised by RNase treatment, providing direct evidence that RNA can function as a structural component of Tau assemblies rather than simply as a passive molecular crowding agent ^39^. Tau inclusions themselves have been shown to be RNA–protein condensates enriched for small RNA species that mislocalise splicing factors such as SRRM2 from nuclear speckles to the cytoplasm, resulting in dysregulated pre-mRNA splicing^40^. Moreover, Tau itself has been shown to undergo liquid–liquid phase separation at physiological concentrations, promoting the formation of toxic oligomeric species without obvious filament formation, and this is enhanced by disease relevant modifications such as AT8 phosphorylation ^41^. Importantly, stress granule–associated RBPs have been shown to be key modulators of this process. Interaction of Tau with TIA1 drives phase separation and directly promotes highly neurotoxic tau oligomers, whereas loss of TIA1 attenuates tau-dependent neurodegeneration in vivo ^42^. Conversely, G3BP2 acts as a protective factor that prevents Tau aggregation, with depletion of G3BP2 in ALS iPSC-MNs markedly increasing tau pathology ^43^. These context-specific effects suggest that Tau toxicity is not determined simply by whether Tau aggregates, but by the material state, reversibility and molecular composition of Tau-associated assemblies. Indeed, recent work has shown that TDP-43 and Tau directly co-condense, with TDP-43 suppressing Tau fibril formation and Tau seeding while promoting small oligomeric Tau/TDP-43 assemblies and TDP-43 seeding activity ^44^. Thus, suppression of Tau fibrillisation can coincide with a shift towards alternative assembly states that may remain biologically deleterious. Further work will be required to define how exon 10-containing MAPT transcripts influence these properties in cellular contexts, to determine whether they directly modulate Tau phase behaviour or act indirectly through RNA-binding proteins, and to establish how these mechanisms contribute to neuronal vulnerability in vivo.

An interesting finding of this study was that exon 10-containing MAPT transcripts appeared to promote the resolubilisation of Tau aggregates. This contrasts with previous reports of RNA–Tau interactions, in which RNA more commonly promotes Tau condensation or fibrillisation ^34,39^, a behaviour more consistent with the non-exon 10-containing transcripts examined here. However, aggregation should not be considered inherently pathogenic. In some contexts, the formation of larger assemblies may represent a protective sequestration mechanism that limits the availability of more toxic soluble species, as has been proposed for stress granules and inclusion bodies and that the soluble oligomeric species can be more pathogenic ^45,46^. Therefore, exon 10-containing MAPT RNA may not simply prevent Tau aggregation, but instead remodel Tau phase behaviour towards a dynamic soluble oligomeric state with greater pathogenic potential.

One potential mechanism is through sequestration or dysregulation of RBPs that aberrantly bind to exon 10. Both FUS and SFPQ are known to regulate alternative splicing and RNA localisation and the increased association of SFPQ with *MAPT* exon 10 in the cytoplasm of *VCP^mu^* neurons may contribute to feed-forward mis-splicing or dysfunction of other critical transcripts. The toxicity of RNA species, independent of their protein products, is increasingly recognised in neurodegeneration, with precedent from repeat expansion disorders such as C9orf72-ALS ^47^, myotonic dystrophy, and fragile X-associated tremor/ataxia syndrome. In this context, the *4R MAPT* RNA may function similarly as an RNA-based toxic species, potentially via the formation of abnormal RNA–RBP complexes or by overwhelming the splicing machinery. The absence of neuroprotection from total *MAPT* knockdown, in contrast to the rescue from 4R-specific siRNA, supports a model where RNA species derived from exon 10 inclusion are the key mediators of toxicity.

This work also fits within a broader view of neuronal transcripts as compartmentalised regulatory molecules rather than passive templates for protein synthesis, with their disruption being a hallmark of ALS ^5^. Previous work in sympathetic neurons demonstrated that axonally localised RNAs can exert coding-independent functions through local RNA–protein interactions^24^, while local processing of axonal transcripts can generate functionally distinct RNA species required for axon integrity^48^. More broadly, long mRNAs have recently been shown to act as signalling scaffolds capable of organising protein networks within defined subcellular compartments^25^. This is particularly relevant for MAPT, whose transcript contains established localisation information within its 3′UTR and is subject to regulated subcellular targeting in neurons ^49^. Although exon 10 itself is not known to act as an axonal localisation element, its inclusion may alter MAPT RNA structure or RBP occupancy in a way that modifies how existing localisation elements are interpreted. In the context of ALS-associated nuclear depletion and cytoplasmic accumulation of FUS, SFPQ and TDP-43, aberrant exon 10 inclusion may therefore perturb not only *MAPT* splicing, but also the localisation, assembly or local function of *MAPT*-containing RNPs.

Some noteworthy limitations include that we did not directly assess axonal localisation or local translation of exon 10-containing MAPT transcripts, and future compartment-specific imaging or axon-enriched RNA profiling will be required to test this model. Furthermore, RNA folding was predicted using isolated exon fragments rather than the full-length MAPT transcript; although these models cannot fully recapitulate native RNA conformation, they nonetheless demonstrate that exon 10 inclusion intrinsically favours a more structured, self-associating conformation, consistent with our experimental data. Additionally, 4R Tau protein remained below the detection threshold of our assays, so we cannot formally exclude a contribution from low-abundance or locally translated 4R Tau; however, the rapid kinetics of the survival phenotype relative to Tau’s known half-life strongly favour an RNA-driven mechanism. Finally, our RNA aggregation and Tau condensation assays use simplified recombinant systems that do not capture the full complexity of the neuronal cytoplasm; the loss of Tau condensates upon addition of exon 9-10 RNA indicates altered phase behaviour, though the fate and material state of the resulting Tau pool remains to be defined. Resolving these points in cellular and in vivo contexts is an important direction for future work.

In conclusion, we propose a model in which RBP mislocalisation disrupts normal splicing regulation of *MAPT*, leading to cytoplasmic accumulation of 4R *MAPT* RNA and increased neuronal vulnerability. These findings suggest that targeting Tau isoform expression or its RNA-binding interactions may offer a therapeutic strategy in ALS, even in the absence of overt tau protein aggregation.

**Extended Data Figure 1:**
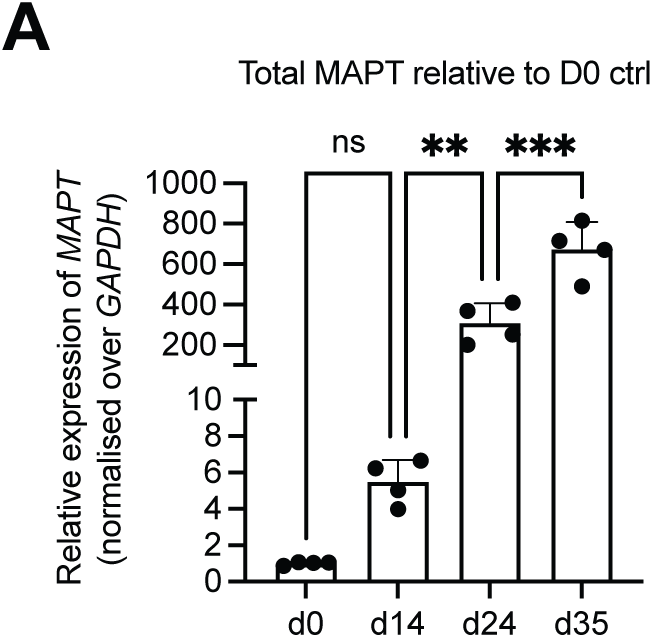
**A** qRT-PCR analysis of expression of Total *MAPT* in CTRL iPSC (d0), motor neuron precursors (iPSC-MN precursors; d14) and motor neurons, (iPSC-MNs; d24, d35) Values are normalised over *GAPDH* and expressed as fold change over D0. Data points indicate individual lines. Please note the break in the Y axis for visibility of data. One way ANOVA with Tukeys with significance placed at P<0.05. ns = not significant, * =P<0.05 **=P<0.001 ***=P<0.0001

**Extended Data Figure 2:**
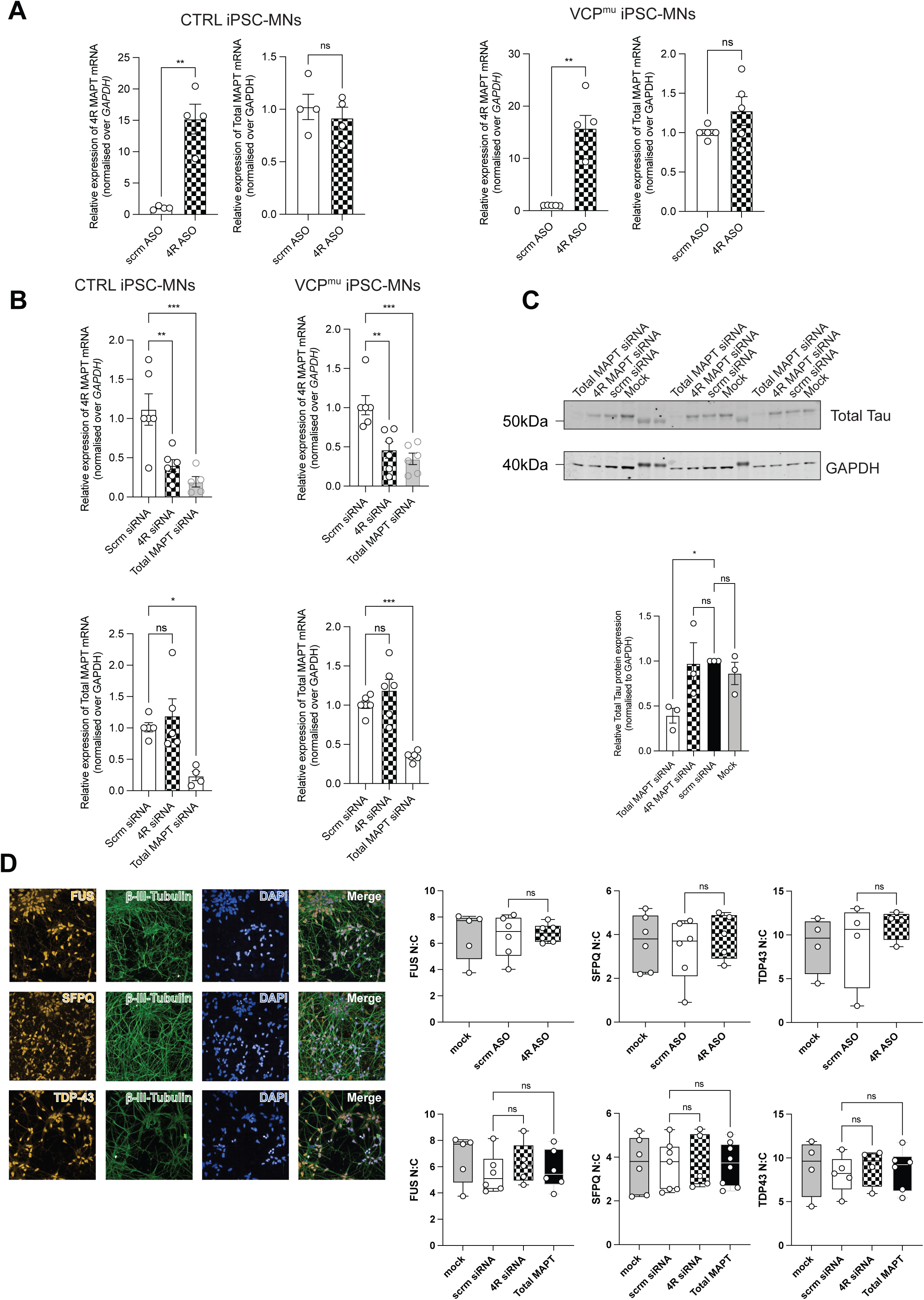
**A** iPSC-MNs were transfected with splice-switching antisense oligonucleotide (ASO) promoting exon 10 inclusion (4R ASO) ^30^ or scrambled control. Expression of exon 10 containing transcripts (left) or total *MAPT* transcripts (right) quantified by qRT-PCR and normalised over *GAPDH* and displayed as fold change over scrambled control. Data points indicate individual lines. Unpaired t-Test with Welch’s correction with significance placed at P<0.05. **B.** As for (A) but utilising an siRNA targeting *MAPT* exon 10 selectively (*4R MAPT*), all *MAPT* isoforms (total *MAPT)* or a scrambled control. Data points indicate individual lines. One way ANOVA with significance placed at P<0.05. **C (**Left) Representative western blots showing total Tau, and GAPDH loading control in iPSC-MNs transfected with an siRNA targeting *MAPT* exon 10 selectively (*4R MAPT*), all *MAPT* isoforms (total *MAPT)* or a scrambled control. (Right). Densitometry of blots. Values are normalised over GAPDH levels and expressed as fold change over scrambled control. Data points indicate individual lines. One way ANOVA with significance placed at P<0.05. **D** (Left) Representative immunofluorescence images of control iPSC-MN stained with FUS, SFPQ or TDP-43 (red) with DAPI (blue) and βIII-tubulin (green) marking nuclei and neurites, respectively. (Right) Quantification of N:C ratios of FUS (Left) SFPQ (centre) and TDP-43 (right), in CTRL iPSC-MNs transfected as indicated. Data represent mean ± SEM across three independent CTRL from two independent experimental repeats. One way ANOVA with significance placed at P<0.05. ns = not significant, * =P<0.05 **=P<0.001 ***=P<0.0001

**Extended Data Figure 3.**
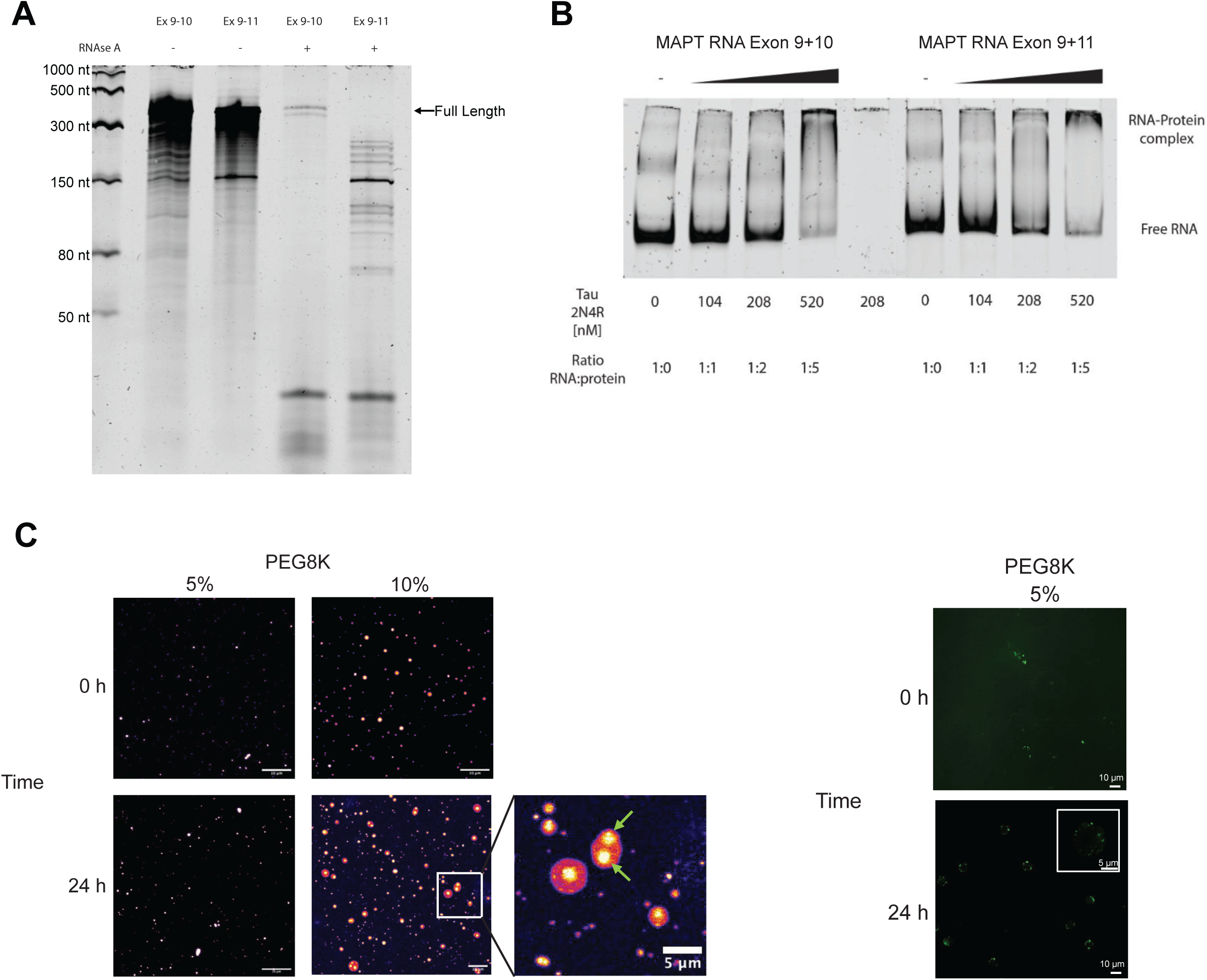
**A** Representative denaturing urea-PAGE analysis of in vitro-transcribed MAPT exon 9-10 and exon 9-11 RNA following treatment with RNase A (50 ng/mL). RNA was incubated at 5 μM under near-physiological buffer conditions (10 mM HEPES, 50 mM NaCl, 2 mM DTT), followed by RNase A digestion for 30 min at room temperature. Reactions were terminated by Proteinase K treatment prior to denaturation and separation on a 7% urea-polyacrylamide gel. RNA was visualised using SYBR Gold staining. Exon 9-10 RNA retained a greater proportion of full-length transcript following RNase A digestion than exon 9-11 RNA, consistent with reduced accessibility to single-stranded RNA cleavage. **B** Representative electrophoretic mobility shift assay (EMSA) using in vitro-transcribed MAPT exon 9-10 or exon 9-11 RNA incubated with increasing concentrations of recombinant human 2N4R Tau protein (0, 104, 208 and 520 nM). RNA was maintained at a constant concentration of 104 nM, generating Tau molar ratios of 0:1, 1:1, 1:2 and 1:5. Formation of RNA-protein complexes was evident as a mobility shift with a corresponding reduction in free RNA. **C.** Representative fluorescence micrographs showing *in vitro* aggregation of recombinant (*Left*) 4R Tau and (*Right*) 3R Tau (2.5 µM). Reactions were performed in a buffer containing 10 mM HEPES, 50 mM NaCl, and 2 mM DTT at room temperature, with PEG-8K added at the indicated concentrations (5% or 10%). Samples were stained with SYPRO Orange to visualise protein aggregates. Images show aggregates at 0 h and 24 h after incubation. Increasing PEG-8K concentration promotes the formation of larger and more numerous Tau aggregates over time, consistent with crowding-induced protein condensation and aggregation. The boxed region highlights representative aggregates, shown enlarged at right. Arrowheads indicate regions of increased fluorescence intensity within aggregate cores, suggesting local enrichment or densification of Tau protein. Scale bars: 10 µm (main panels) and 5 µm (zoomed inset).

## Methods

### Ethics statement

Informed consent was obtained from all patients and healthy controls who donated samples for hiPSC culture. Experimental protocols were conducted according to approved regulations and guidelines by UCLH’s National Hospital for Neurology and Neurosurgery and UCL’s Institute of Neurology joint research ethics committee (09/0272).

### Human iPSC-derived MN lines

iPSC lines were derived from skin biopsies of healthy donors, sporadic ALS patients or ALS patients harboring VCP mutations. Informed consent was obtained from all subjects. Experimental protocols were all carried out according to approved regulations and guidelines by UCL Hospitals National Hospital for Neurology and Neurosurgery and UCL Institute of Neurology joint research ethics committee (09/0272). Details of all lines used in this study are found in Table S1. iPSCs were maintained with Essential 8 Medium media (Life Technologies) on Geltrex (Life Technologies) at 37°C and 5% carbon dioxide. iPSCs were passaged when reaching 70% confluency using EDTA (Life Technologies, 0.5 mM). iPSC cultures underwent differentiation into spinal cord motor neurons as previously described^28^ Briefly, iPSCs were plated to 100% confluency and then differentiated to a spinal neural precursor fate by sequential treatment with small molecules, 0–7: 1 μM Dorsomorphin (Tocris Bioscience), 2 μM SB431542 (Tocris Bioscience), and 3.3 μM CHIR99021 (Tocris Bioscience), day 7–14: 0.5 μM retinoic acid (Sigma Aldrich) and 1 μM Purmorphamine (Sigma Aldrich), day 14–18: 0.1 μM Purmorphamine. After neural conversion and patterning, 0.1 μM Compound E (Bio-techne) was added to promote terminal differentiation into spinal cord motor neurons.

### RT-qPCR

RNA was extracted using the Maxwell RSC simplyCells RNA kit (Promega) and the Maxwell RSC instrument (Promega), according to the manufacturer’s instructions. Contaminating genomic DNA was removed by DNAse I digestion during the RNA extraction. RNA concentration and purity ratios were determined by NanoDrop (ThermoFisher Scientific). RevertAid First Strand cDNA Synthesis kit (ThermoFisher Scientific) or SuperScript IV (ThermoFisher Scientific) was used to synthesise cDNA. Typically, 200-500 ng RNA was reverse transcribed per each reaction that employed random hexamers. Appropriate dilution of the cDNA was then used in qPCR reactions containing PowerUp SYBR Green Master Mix and primer pairs, performed in triplicate using the QuantStudio 6 Flex Real-Time PCR System (Applied Biosystems). Reactions omitting reverse transcriptase and containing no template were used as controls. Primers were determined to have 90-110% efficiency. Primers used in this study are listed in Table S3. Data were analyzed according to the ddCt method, typically expressed as FC over the control or untreated group.

### Protein Extraction and Western Blotting

Proteins were extracted using boiling LDS buffer with 5% beta-mercaptoethanol followed by homogenisation using 25G needle. Protein samples were then denatured for 5 minutes at 95 °C, loaded onto an Invitrogen NuPAGE 4-12% Bis-Tris protein gel (ThermoFisher Scientific) and run at 180 V for 1-2 hours in NuPAGE MOPS running buffer (ThermoFisher Scientific). PageRuler™ Prestained Protein Ladder, 10 to 180 kDa (ThermoFisher Scientific, 26616) was run alongside as the marker of protein molecular weight. Proteins were blotted onto 0.45 µm nitrocellulose membrane using XCell II™ Blot Module wet transfer apparatus at 330 mA for 2 hours. Membranes were blocked in TBS containing 0.1% Tween-20 (TBST) and 5% BSA for one hour at room temperature. Incubation with an appropriate dilution of the primary antibody in BSA was carried out overnight at 4 °C. Three 10-minute washes were then performed in TBST, followed by incubation with an appropriate fluorescent IRDye® 680 or 800CW secondary antibody (LI-COR Biosciences) at 1:10,000 dilution in TBST, for one hour at room temperature. After a further three 10-minute washes in TBST, the signal was imaged using Odyssey® CLx Infrared Imaging System (LI-COR Biosciences). Antibodies used in this study are found in Table S2

### ASO design

Splice-switching antisense oligonucleotides (ASOs) were designed with a phosphorothioate backbone and had the following modifications on a phosphorothioate (PS) backbone: uniformly modified with 2′-O-methoxyethyl nucleotides to improve nuclease resistance and prevent RNaseH-mediated degradation of *MAPT* mRNA. The 3R to 4R MAPT ASO sequence was GGCGCATGGGACGTGTGA as described previously ^30^.

### siRNA design

siRNAs targeting human total MAPT (pool of 4 siRNAs, ON-TARGETplus Human MAPT siRNA, L-012488-00-0005) and isoforms containing exon 10 were obtained from Horizon. The Exon 10 specific siRNA was designed in collaboration with Horizon bioscience and the sequence used was TATCCTTTGAGCCACACTT. As a negative control the ON-TARGET*plus* Non-targeting Pool (Horizon D-001810-10) was used.

### Transfection

Cells were transfected using Lipofectamine RNAiMAX (ThermoFisher Scientific), according to manufacturer’s instructions, with 100 nM of appropriate siRNA and 50 nM of ASO used per transfection. Scramble siRNA or ASO was used as a negative control while the mock sample was treated with the empty transfection reagent.

### Cell survival analysis

For live imaging to assess neuronal survival, neural precursors were dissociated using Accutase (Life Technologies) and plated into 96-well plates (Falcon) coated with polyethylenimine (PEI; 2.2 mg/ml in water; Sigma) and Geltrex. Cells were transfected as detailed above before media was changed into fresh N2B27 containing compound E and 1:200 NucView Caspase-3 dye (Sartorius) and 1:1500 Nuclight Red nuclear dye (Sartorius) and imaged using the 10 × objective of an Incucyte imaging platform with five fields per well condition. Images were acquired every 3h for 6 days total.

### Immunocytochemistry

Neural precursors were dissociated using Accutase (Life Technologies) and plated into 96-well plates (Falcon) coated with polyethylenimine (PEI; 2.2 mg/ml in water; Sigma) and Geltrex and transfected as described above. Following 24 h, media was changed to N2B27 + Compound E and cells differentiated for 6 days. Media was removed and cells were washed once in PBS prior to fixing in 4% paraformaldehyde in PBS for 10 minutes at room temperature (RT). Cells were washed twice in PBS and were permeabilised and non-specific antibody binding was blocked using 5% Bovine Serum Albumin (BSA) (Sigma-Aldrich, A7030) diluted in PBS containing 0.3% Triton X-100 (PBSTx) for 60 minutes at RT. Primary antibodies were prepared in 5% BSA/PBSTx at the required dilution (Table S2) and applied to cells overnight at 4°C. Cells were washed twice in PBS before primary antibody detection using InvitrogenTM Alexa FluorTM secondary antibodies (1:1000) and DAPI nuclear stain (1:2000 in 5% BSA/PBSTx) for 1 hour in the dark at RT. Secondary antibodies and DAPI were removed using 2 further PBS washes. Cells were then imaged as detailed below.

### Intracellular ROS measurement

Oxidative stress was measured by detection of intracellular ROS using a fluorogenic probe, CellROX™ Green (ThermoFisher). neural precursors were dissociated using Accutase (Life Technologies) and plated into 96-well plates (Falcon) coated with polyethylenimine (PEI; 2.2 mg/ml in water; Sigma) and Geltrex and transfected as described above. Following 24 h, media was changed to N2B27 + Compound E and cells differentiated for 48 h. The dye stock + Hoechst solution was prepared in 30 µl maintenance media per well and added to 100 µl of maintenance media already in the well to make up a final concentration of 5 µM, followed by a 30-minute incubation in the dark at 37°C. Cells were then washed once with PBS, before imaging in PBS. Cells were then imaged as described below.

### High throughput confocal imaging and analysis

HiPSC-derived motor neurons plated in 96-well formats were visualised using the Perkin Elmer Opera Phenix™ High Content Screening System with a 20x objective. For each well a minimum of 8 fields were acquired. Z stacks of images were acquired with a minimum of 10 slices per stack, with images displayed as maximum projections. Acquisition and thresholding settings were standardised for each experimental block. Cells were analysed with the complementary Columbus™ Image Data Storage and Analysis system.

### Cell Fractionation

Biochemical subcellular fractionation was performed as described previously ^35^. Briefly, samples were processed using the Ambion PARIS kit (ThermoFisher Scientific) cell fractionation buffer, following the manufacturer’s general protocol, and an 8 M Urea Nuclear Lysis Buffer. Cells were washed once using ice-cold PBS. Cytosolic fraction was then obtained by lysing cells directly in ice-cold cell fractionation buffer (ThermoFisher Scientific) for 3 minutes, Lysates were then centrifuged for 3 minutes at 500 x g at 4°C. The supernatant was then collected, further centrifuged at maximum speed in a bench centrifuge at 4°C for 1 minute, and the resulting supernatant was then processed as cytosolic fraction. Nuclear pellets from the first centrifugation step were gently washed once with cell fractionation buffer and then lysed on ice for 30 minutes in 8 M Urea Nuclear Lysis Buffer, containing 50 mM Tris-HCL (pH 8), 100 mM NaCl, 0.1% SDS, and 1 mM DTT. The resulting nuclear fraction was then homogenised using a QIAshredder (QIAGEN) to shred chromatin and reduce viscosity, before being further processed for RNA extraction. Both lysis buffers were supplemented with 0.1 U/µl RiboLock RNase Inhibitor (ThermoFisher Scientific) and HALT Protease Inhibitor Complex (ThermoFisher Scientific).

### iiCLIP Improved individual-nucleotide resolution UV-crosslinking and immunoprecipitation of protein-RNA complexes (iiCLIP)

To identify SFPQ RNA-binding sites in hiPSC-derived MNs, iiCLIP^50^ was performed at DIV24 of neuronal differentiation. Cells at 90% confluence were uniformly irradiated with UV-C on ice (150 mJ/cm²; Stratalinker 2400, 254 nm) to covalently cross-link RBP–RNA interactions. Cells were then lysed in lysis buffer containing Complete protease inhibitor cocktail (Roche, 25×). RNA was fragmented by adding 0.1 U RNase I and 4 U Turbo DNase per 1 mL of cell lysate at 1 mg/mL protein concentration. A high-RNase control was treated with 2.5 U RNase I.

RBP–RNA complexes were purified using antibodies against human SFPQ (mouse anti-SFPQ, Abcam, Cat# ab118525, clone B92) for SFPQ-iiCLIP. Antibodies were coupled to magnetic Protein G Dynabeads (Cat#10004D) to isolate protein–RNA complexes. For complex visualisation and sample multiplexing, RNA was ligated to pre-adenylated, IR-labelled barcoded L7 adapters (sequences in Table S3). Complexes were size-separated by SDS-PAGE, transferred to nitrocellulose, and visualised using an Odyssey CLx Scanner (LI-COR) to assess RNA digestion and confirm the expected RNA–protein complex size. RNA was released by proteinase K digestion and recovered by precipitation. RNA fragments were reverse-transcribed using Superscript IV Reverse Transcriptase (Life Technologies, #18090010), purified with AMPure XP beads (Beckman-Coulter, Cat#A63880), and circularised using Circligase II (Epicenter, #CL9021K), followed by AMPure XP purification. Libraries were PCR-amplified and size-selected using AMPure beads, then sequenced as single-end 100-bp reads on an Illumina HiSeq 4000 or NovaSeq.

### iiCLIP data processing and computational analysis

Sequencing reads were demultiplexed using Ultraplex^51^ with the adapter sequence AGATCGGAAGAGC and allowing one mismatch in the 5′ barcode. Reads were processed using the standardised iiCLIP workflow implemented in the nf-core/clipseq Nextflow pipeline v1.0.0 (https://nf-co.re/clipseq). The human GRCh38 genome and GENCODE v38 annotation were used as references. Downstream analyses were performed using custom R scripts (v4.1.3).

iiCLIP crosslink counts for SFPQ were normalised to experimental library size and smoothed across regions of interest using a 25-nt rolling mean. Smoothed, normalised values were plotted across the regions. Trackplot v0.5.3 (https://trackplot.readthedocs.io/en/latest/) was used to visualise RNA-seq coverage and iiCLIP tracks. RNA-seq BAM files generated using STAR/Salmon within nf-core/rnaseq v1.0.0 and iiCLIP peak bigWig files generated using clipplotr v1.0.0 (https://github.com/ulelab/clipplotr) were imported into Trackplot or clipplotrviz to visualise expression and binding signals as crosslink or read coverage counts per million (CPM).

### RNA immunoprecipitation

RNA immunoprecipitations were carried out as previously described ^24^ with the following modifications. Three μg SFPQ antibody (Abcam, ab11825), or Mouse IgG antibody (Santa Cruz sc2027) were incubated with prewashed protein G dynabeads (Thermo) in a PBS buffer containing 1 mg/ml heparin and 1% BSA for 2 hours 4°C, and then washed with wash buffer (50 mM Tris pH8, 150 mM NaCl, 1% Triton X-100). 100 μg of cytoplasmic lysate from control or VCPmu MN samples was incubated with antibody conjugated beads for 1 hour at 4°C in wash buffer containing 0.2 mg/ml heparin, 0.1 U/μl RiboLock RNase Inhibitor (Thermo) and Halt protease inhibitor cocktail. 2% of lysate volume was purified by the Maxwell RSC simplyCells kit (Promega) for total input. Beads were washed 6 times with wash buffer 5 minutes 4°C followed by elution of RNA in extraction buffer (0.2 M NaAcetate, 1 mM EDTA, 0.2% SDS) for 5 minutes at 70°C, or protein in 2x LDS 5% beta mercaptoethanol and boiled for 8 minutes at 95°C. RNA was purified from immunocomplexes using PureLink® RNA Micro Scale Kit according to manufacturer’s instructions with on-column DNAse digestion, reverse transcribed with Superscript IV and random hexamers (Thermo) and then analyzed by qPCR.

### RNA structure prediction

We used the ViennaRNA Package^52^ to calculate the secondary structure of the RNA sequence corresponding to exon 9, 10, 11 and their combination. We used the RNAfold Web server interface and calculated Minimum Free Energy and partition function at 37C using RNA parameters from Turner model (2004).

Since ViennaFold can predict only unimolecular folding of RNA, we tested the sequence for multimerization using NUPACK using rna95 parameter set and a concentration of 5 µM .

### In vitro RNA transcription

Templates for in vitro transcription were generated by PCR to append a T7 promoter upstream of the Tau exons sequences. For the exon 9 + exon 10 template, the region was amplified directly from the plasmid pRK5-EGFP-Tau_1 (Addgene #46904) using primers T7-exon9+10_Fw and T7-exon9+10_Rev (Table S3), which introduced the T7 promoter sequence. For the exon 9 + exon 11 template, pRK5-EGFP-Tau_1 was first subjected to site-directed mutagenesis with Q5 Site-Directed Mutagenesis Kit (New England Biolabs), to remove the exon 10 coding sequence, using primers Exon9+11-Mut_Fw and Exon9+11-Mut_Rev according to manufacturer’s instructions. The resulting construct was then amplified with primers T7-exon9+11_Fw andT7-exon9+11_Rev to insert the T7 promoter. Both template amplifications were performed with Q5 High-Fidelity 2X Master Mix (New England Biolabs) according to the manufacturer’s instructions. In vitro transcription was carried out using 760 ng of purified template as input for the T7 FlashScribe Transcription reaction (CELLSCRIPT) carried out according to the manufacturer’s instructions. The transcribed RNA was then purified using Monarch Spin RNA Cleanup Kit (New England Biolabs). The length and integrity of the resulting transcripts were confirmed on a denaturing PAGE gel.

### RNAse A digestion

RNA sequences were synthesised by *IVT* as described above and diluted to a final concentration of 5 µM in assay buffer (HEPES 10mM, NaCl 50mM, DTT 2mM). RNase A (ThermoFisher) was added to a final concentration of 50 ng/mL. Samples were incubated at room temperature (22 C) for 30 min, after which Proteinase K (ThermoFisher, 1 U/mL) was added to the sample to remove degrade protein components previous to gel loading. Samples were diluted 1:10 with assay buffer, combined with RNA loading dye (NEB) and denatured at 95 °C for 1min prior to quench on ice. Samples were run on a polyacrylamide denaturing gel (6% acrylamide, 7M urea) for 45 min at 120 V in TBE 1x. After the run, gel was stained with SYBR Gold (ThermoFisher, 1x) and imaged on a Li-Cor Odyssey Imaging System in the SYBR Gold excitation/emission channel.

### EMSA assay

RNA and Tau (2N4R) protein (HY-P70589 MedChemExpress) working stocks were prepared by dilution in nuclease-free water to 2 µM. Binding reactions (12 µL) were assembled on ice and contained 1× binding buffer (25 mM HEPES pH 7.2, 100 mM KCl, 2 mM MgCl₂, 1 mM DTT, 10% (v/v) glycerol), 0.1 mg/mL BSA, and (0.2 U/µL) RNase inhibitor, together with 40 nM RNA (exon 9 + 10 or exon 9 + 11) and Tau protein at the indicated concentrations (0–520 nM); nuclease-free water was added to the final volume. Reactions were incubated at room temperature for 30 min and complexes were resolved on a 6% native polyacrylamide gel run at 90 V for 180 min at 4 degrees. Gels were stained with SYBR Gold (ThermoFisher) for 25 min and visualized on a Li-Cor Odyssey Imaging System.

### RNA Aggregation in Mg^2+^ buffer

Sample preparation. RNA of interest from a stock solution in water is diluted to the desired concentration in a buffer prepared by mixing RNase free water, HEPES (final concentration 10mM), NaCl (final concentration 25mM) and MgCl2 (final concentration 50mM). No crowding agent was used in these experiments. The samples are incubated at room temperature for 30 min, before adding SYBR Gold (0.5x final concentration). Glass coverslips (1.5H, 0.170mm) were previously cleaned with Hellmanex III 2% and extensively washed with water and ethanol. An imaging spacer (GraceBio, 9mm, 0.12 mm thickness) was applied to the coverslip, 5 µL of samples were added and sealed with a second coverslip applied on top. Samples were imaged on a Leica SP8 using a 10x air objective with zoom 2x, using SYBR Gold settings available on LAS X software. Images were processed using Li Autothresholding method, followed by Fiji’s Analyze Particle to measure the area of assemblies per FoV. For each FoV, we calculated the ratio between the total area of the assembly over the FoV area. Each experimental conditions was performed in duplicate using independent stock of RNA (n=2), for each sample three FoV were acquired (N=3).

### RNA and protein condensation

*Sample preparation.* RNA of interest was diluted to the desired concentration in freshly prepared and filtered assay buffer (HEPES 10mM, NaCl 50mM, DTT 2mM). Tau (2N4R HY-P70589 MedChemExpress) was then added to the solution and ultimately PEG 8K (Sigma-Aldrich) from a freshly prepared 40% w/v stock solution was added to the system to promote phase separation. Samples were incubated at room temperature for 30 min, SYPRO Orange 1x (ThermoFisher) was added to allow visualization of protein condensates. Glass coverslips (1.5H, 0.170mm) were previously cleaned with Hellmanex III 2% and extensively washed with water and ethanol. An imaging spacer (GraceBio, 9mm, 0.12 mm thickness) was applied to the coverslip, 5 µL of samples were added and sealed with a second coverslip applied on top. Samples were kept in the dark until imaging. Imaging was performed on a Leica SP8 confocal microscope using 60x oil immersion objective, excitation wavelength set at 458nm and emission between 557-650 nm (HyD detector). Each experimental condition was performed in duplicate (n=2) and for each sample at least four fields of view (FoV, N=4) spanning different areas of the samples were recorded. For each FoV, z-stack of at least 15 µm were acquired. Maximum intensity projection of each z-stack was thresholded using Otsu thresholding algorithm and droplets area measured with Fiji (ImageJ) “Analyze particle” plugin. Droplets with circularity below 0.85 and area below 0.5 µm^2^ were excluded from the analysis.

### Effect on disease progression in postmortem ALS samples

We processed postmortem RNA-seq samples for ALS, ALS+FTD and control patient groups from the NYGC-ALS consortium project PRJNA644618 in GEO. In total, we analysed 2256 polyA -ve RNA-seq samples mapping to 1837 tissue samples. Raw FASTQs were aligned to the GRCh38 reference genome using STAR aligner. We used additional alignment parameters “*--outSAMattributes All*” and “*--outSAMtype BAM SortedByCoordinate”*. In total, we discarded 197 samples with mapping percentage less than 90%. We further removed 340 samples containing an invalid RIN score (RIN score <= 4 or NA; 137) or containing samples from other neurodegenerative disorders (203). For downstream analyses we used 1300 BAM files which passed all previous filtering steps.

We used the *featureCounts* function within Rsubread package in R/Bioconductor to quantify fractional counts within non-overlapping exonic and intronic features identified in ENSEMBL transcriptome annotation v104 using *exonicParts* and *intronicParts* function from GenomicFeatures package in R/Bioconductor. BAM files were provided as input through the “*files*” argument. Features were provided as an input through the *annot.ext* argument and setting the “*useMetaFeatures=T”*. We set “minMQS=255” with 255 being the minimum mapping quality set by STAR aligner for uniquely mapping reads. To obtain fractional counts we set *“fraction=T”* and *“allowMultiOverlap=T”*. Additionally, We set *“isPairedEnd”, “strandSpecific”, and “requireBothEndsMapped”*.

We obtained the total number of fractional counts mapping to *MAPT E10* (chr17:46010306:46010402:+). For each patient, we aggregate multiple samples from the same sub-anatomical region to its mean. Separately, we obtained the total number of genic counts (exons) for *MAPT*. We computed the E10 usage as MAPT E10 CPM divided by the MAPT CPM.

We used the *randomForestSRC* package in R and implemented Random survival forests using the *rfsrc* function. To control for variation due to starting seeds, patient dichotomisation is performed using the median RSF risk score. The median risk score is obtained from a distribution of RSF risk scores derived from 100 RSF models generated using 100 unique starting seeds. Furthermore, we used additional parameters *ntree = 1000, importance = TRUE*.

For investigating patient progression in postmortem ALS cervical spinal cord samples, MAPT expression in CPM units. We used the disease duration as the time parameter and the MAPT expression to construct a survival function, including age of onset and genotypic sex as covariates. We used this survival function as an input for the RSF function and investigated the effect of MAPT expression as covariates. The RSF risk score was predicted using the *predict* function.

### Statistical Analysis

Statistical analysis and associated graph generation were conducted using GraphPad Prism. Statistical tests used for each experimental dataset are reported within the figure legends, when significant differences were identified, alongside repeat numbers. Data are presented as means ± SEM. Statistical significance was accepted at a P value of <0.05 and either displayed as actual value or star (*P ≤ 0.05, **P ≤ 0.01, ***P ≤ 0.001, or ****P ≤ 0.0001).

## Data / code availability

Further information and requests for resources should be directed to the lead contacts, Rickie Patani, Marco di Antonio and Hamish Crerar

## Acknowledgements

We thank Selina Wray and her laboratory for the generous donation of patient-derived fibroblasts and iPSC lines, and we are especially grateful to the individuals who kindly donated skin biopsies for the derivation of ALS and control hiPSC lines used in this study. We thank all members of the Patani, Ruepp and diAntonio labs for their valuable feedback and technical support. We kindly thank Christy Hung for her support in providing the ASOs for 3R-4R splice site switching and controls. We gratefully acknowledge the Francis Crick Institute’s High Throughput Screening Science Technology Platform for imaging support. H.C. gratefully acknowledges funding from a My Name’5 Doddie Foundation catalyst award (MN5DF/CAAu23/100027) and a Motor Neuron Disease Association non-clinical junior fellowship (Crerar/Oct21/974-799). R.P. gratefully acknowledges generous support from a Lister Research Prize Fellowship, Steve Redgwell, Liane Iles, Challenging MND, the Motor Neuron Disease Association (Patani/Dec22/957-793), My Name’5 Doddie Foundation (MN5DF/2022/003), and Target ALS (BB-2024-C4-L4). This work was also supported by the Francis Crick Institute, which receives its core funding from Cancer Research UK, the UK Medical Research Council, and the Wellcome Trust, the KCL Dementia Research Institute which receives its core funding from the Medical Research council, and M.D.A acknowledges support from Lister Research Prize Fellowship 2022. M.S. is supported by the European Union’s Horizon 2020 research and innovation program under the Marie Skłodowska-Curie grant agreement No. 101203352 (MiMoSeS). M.S. and M.D.A. acknowledge the support of Imperial College London, Leica Microsystem Imaging Hub (at Bioengineering Department) for providing access to microscopy instrumentation. Paper proofreading was assisted by ChatGPT 5 (OpenAI) and Claude Opus 4.8 (Anthropic).

## Author Contributions

Conceptualization was done by H.C. and R.P.

Experiments were performed by H.C. M.S. S.P.N. M.P.H. G.M. Y.W., B.C.

Incucyte longitudinal cell death imaging and analysis: H.C. and B.C.

RBP mislocalisation imaging and analysis: H.C. and Y.W.

Oxidative stress analysis: H.C. iiClIP and analysis: G.M.

RNA Immunoprecipitation: H.C.

RNA, protein aggregation and *in vitro* assays: M.S. and S.P.N.

Postmortem transcriptomic analysis: K.P.

Data analysis was done by H.C., M.S., S.P.N., G.M. and K.P.

Resources were provided by M.P.H, Y.W and B.L.-C.

Supervision: R.P., M.d.A., M-D.R.

Writing of the original draft was done by H.C. and R.P.

Review and editing were done by R.P, H.C, M.dA, M.S and M-D.R.

Funding acquisition for the work in this study was done by R.P. and H.C.

## References

1. Neumann, M. et al. Ubiquitinated TDP-43 in frontotemporal lobar degeneration and amyotrophic lateral sclerosis. Science 314, 130–133 (2006).

2. Tyzack, G. E. et al. Widespread FUS mislocalization is a molecular hallmark of amyotrophic lateral sclerosis. Brain 142, 2572–2580 (2019).

3. Luisier, R. et al. Intron retention and nuclear loss of SFPQ are molecular hallmarks of ALS. Nat. Commun. 9, 2010 (2018).

4. Diaz-Garcia, S. et al. Nuclear depletion of RNA-binding protein ELAVL3 (HuC) in sporadic and familial amyotrophic lateral sclerosis. Acta Neuropathol 142, 985–1001 (2021).

5. Ziff, O. J. et al. Nucleocytoplasmic mRNA redistribution accompanies RNA binding protein mislocalization in ALS motor neurons and is restored by VCP ATPase inhibition. Neuron 111, 3011–3027.e7 (2023).

6. Bryce-Smith, S. et al. TDP-43 loss induces cryptic polyadenylation in ALS/FTD. Nat. Neurosci. 1–11 (2025).

7. Gordon, P. M., Hamid, F., Makeyev, E. V. & Houart, C. A conserved role for the ALS-linked splicing factor SFPQ in repression of pathogenic cryptic last exons. Nat. Commun. 12, 1918 (2021).

8. Brown, A.-L. et al. TDP-43 loss and ALS-risk SNPs drive mis-splicing and depletion of UNC13A. Nature 603, 131–137 (2022).

9. Neeves, J. et al. An alternative cytoplasmic SFPQ isoform with reduced phase separation potential is up-regulated in ALS. Sci. Adv. 11, eadt4814 (2025).

10. Melamed, Z. ‘ev et al. Premature polyadenylation-mediated loss of stathmin-2 is a hallmark of TDP-43-dependent neurodegeneration. Nat Neurosci 22, 180–190 (2019).

11. Klim, J. R. et al. ALS-implicated protein TDP-43 sustains levels of STMN2, a mediator of motor neuron growth and repair. Nat Neurosci 22, 167–179 (2019).

12. Ma, X. R. et al. TDP-43 represses cryptic exon inclusion in the FTD-ALS gene UNC13A. Nature 603, 124–130 (2022).

13. Ishigaki, S. et al. Altered Tau Isoform Ratio Caused by Loss of FUS and SFPQ Function Leads to FTLD-like Phenotypes. Cell Rep. 18, 1118–1131 (2017).

14. Gu, J. et al. TDP-43 suppresses tau expression via promoting its mRNA instability. Nucleic Acids Res. 45, 6177–6193 (2017).

15. Goedert, M., Spillantini, M. G., Potier, M. C., Ulrich, J. & Crowther, R. A. Cloning and sequencing of the cDNA encoding an isoform of microtubule-associated protein tau containing four tandem repeats: differential expression of tau protein mRNAs in human brain. EMBO J. 8, 393–399 (1989).

16. Vintilescu, C. R., Afreen, S., Rubino, A. E. & Ferreira, A. The neurotoxic TAU45-230 fragment accumulates in upper and lower motor neurons in amyotrophic lateral sclerosis subjects. Mol. Med. 22, 477–486 (2016).

17. Lang, A. E., Riherd Methner, D. N. & Ferreira, A. Neuronal degeneration, synaptic defects, and behavioral abnormalities in tau₄₅₋₂₃₀ transgenic mice. Neuroscience 275, 322–339 (2014).

18. Yang, W., Sopper, M. M., Leystra-Lantz, C. & Strong, M. J. Microtubule-associated tau protein positive neuronal and glial inclusions in ALS. Neurology 61, 1766–1773 (2003).

19. Yang, W. & Strong, M. J. Widespread neuronal and glial hyperphosphorylated tau deposition in ALS with cognitive impairment. Amyotroph. Lateral Scler. 13, 178–193 (2012).

20. Chatterjee, M. et al. Plasma extracellular vesicle tau and TDP-43 as diagnostic biomarkers in FTD and ALS. Nat. Med. 30, 1771–1783 (2024).

21. Knight, A. C. et al. Head-to-head comparison of tau-PET radioligands for imaging TDP-43 in post-mortem ALS brain. Mol. Imaging Biol. 25, 513–527 (2023).

22. Hung, C. & Patani, R. Elevated 4R tau contributes to endolysosomal dysfunction and neurodegeneration in VCP-related frontotemporal dementia. Brain 10.1093/brain/awad370 (2023) doi:10.1093/brain/awad370.

23. Ishigaki, S. et al. Aberrant interaction between FUS and SFPQ in neurons in a wide range of FTLD spectrum diseases. Brain 143, 2398–2405 (2020).

24. Crerar, H. et al. Regulation of NGF Signaling by an Axonal Untranslated mRNA. Neuron 102, 553–563.e8 (2019).

25. Chen, X., Fansler, M. M., Janjoš, U., Ule, J. & Mayr, C. The FXR1 network acts as a signaling scaffold for actomyosin remodeling. Cell 187, 5048–5063.e25 (2024).

26. Maharana, S. et al. RNA buffers the phase separation behavior of prion-like RNA binding proteins. Science 360, 918–921 (2018).

27. Iovino, M. et al. Early maturation and distinct tau pathology in induced pluripotent stem cell-derived neurons from patients with MAPT mutations. Brain 138, 3345–3359 (2015).

28. Hall, C. E. et al. Progressive Motor Neuron Pathology and the Role of Astrocytes in a Human Stem Cell Model of VCP-Related ALS. Cell Rep. 19, 1739–1749 (2017).

29. Barmada, S. J. et al. Cytoplasmic mislocalization of TDP-43 is toxic to neurons and enhanced by a mutation associated with familial amyotrophic lateral sclerosis. J. Neurosci. 30, 639–649 (2010).

30. Schoch, K. M. et al. Increased 4R-tau induces pathological changes in a human-tau mouse model. Neuron 90, 941–947 (2016).

31. Silva, M. C. et al. Human iPSC-Derived Neuronal Model of Tau-A152T Frontotemporal Dementia Reveals Tau-Mediated Mechanisms of Neuronal Vulnerability. Stem Cell Reports 7, 325–340 (2016).

32. Cosker, K. E., Fenstermacher, S. J., Pazyra-Murphy, M. F., Elliott, H. L. & Segal, R. A. The RNA-binding protein SFPQ orchestrates an RNA regulon to promote axon viability. Nat. Neurosci. 19, 690–696 (2016).

33. Faraway, R. et al. Collective homeostasis of condensation-prone proteins via their mRNAs. Nature 647, 798–808 (2025).

34. Zhang, X. et al. RNA stores tau reversibly in complex coacervates. PLoS Biol. 15, e2002183 (2017).

35. Tyzack, G. E. et al. Aberrant cytoplasmic intron retention is a blueprint for RNA binding protein mislocalization in VCP-related amyotrophic lateral sclerosis. Brain 144, 1985–1993 (2021).

36. Meyer, H., Bug, M. & Bremer, S. Emerging functions of the VCP/p97 AAA-ATPase in the ubiquitin system. Nat. Cell Biol. 14, 117–123 (2012).

37. Buchholz, S. & Zempel, H. The six brain-specific TAU isoforms and their role in Alzheimer’s disease and related neurodegenerative dementia syndromes. Alzheimers. Dement. 20, 3606–3628 (2024).

38. Sato, C. et al. Tau kinetics in neurons and the human central nervous system. Neuron 97, 1284–1298.e7 (2018).

39. Abskharon, R. et al. Structural evidence that RNA contributes to polymorphism of tau amyloid fibrils. iScience 29, 115501 (2026).

40. Lester, E. et al. Tau aggregates are RNA-protein assemblies that mislocalize multiple nuclear speckle components. Neuron 109, 1675–1691.e9 (2021).

41. Kanaan, N. M., Hamel, C., Grabinski, T. & Combs, B. Liquid-liquid phase separation induces pathogenic tau conformations in vitro. Nat. Commun. 11, 2809 (2020).

42. Ash, P. E. A. et al. TIA1 potentiates tau phase separation and promotes generation of toxic oligomeric tau. Proc. Natl. Acad. Sci. U. S. A. 118, e2014188118 (2021).

43. Wang, C. et al. Increased G3BP2-Tau interaction in tauopathies is a natural defense against Tau aggregation. Neuron 111, 2660–2674.e9 (2023).

44. Simonetti, F. et al. Direct interaction between TDP-43 and Tau promotes their co-condensation, while suppressing Tau fibril formation and seeding. EMBO J. 10.1038/s44318-025-00590-2 (2025) doi:10.1038/s44318-025-00590-2.

45. Arrasate, M., Mitra, S., Schweitzer, E. S., Segal, M. R. & Finkbeiner, S. Inclusion body formation reduces levels of mutant huntingtin and the risk of neuronal death. Nature 431, 805–810 (2004).

46. Takahashi, T. et al. Soluble polyglutamine oligomers formed prior to inclusion body formation are cytotoxic. Hum. Mol. Genet. 17, 345–356 (2008).

47. Raguseo, F. et al. The ALS/FTD-related C9orf72 hexanucleotide repeat expansion forms RNA condensates through multimolecular G-quadruplexes. Nat. Commun. 14, 8272 (2023).

48. Andreassi, C. et al. Cytoplasmic cleavage of IMPA1 3’ UTR is necessary for maintaining axon integrity. Cell Rep. 34, 108778 (2021).

49. Aronov, S., Aranda, G., Behar, L. & Ginzburg, I. Axonal tau mRNA localization coincides with tau protein in living neuronal cells and depends on axonal targeting signal. J. Neurosci. 21, 6577–6587 (2001).

50. Hallegger, M. et al. TDP-43 condensation properties specify its RNA-binding and regulatory repertoire. Cell 184, 4680–4696.e22 (2021).

51. Wilkins, O. G., Capitanchik, C., Luscombe, N. M. & Ule, J. Ultraplex: A rapid, flexible, all-in-one fastq demultiplexer. Wellcome Open Res. 6, 141 (2021).

52. Lorenz, R. et al. ViennaRNA Package 2.0. Algorithms Mol. Biol. 6, 26 (2011).

